# Transcriptomic profiling reveals host-specific evolutionary pathways promoting enhanced fitness in the plant pathogen *Ralstonia pseudosolanacearum*

**DOI:** 10.1101/2023.05.12.540487

**Authors:** Rekha Gopalan-Nair, Marie-Françoise Jardinaud, Ludovic Legrand, Céline Lopez-Roques, Olivier Bouchez, Stéphane Genin, Alice Guidot

**Affiliations:** LIPME, Université de Toulouse, INRAE, CNRS, Castanet-Tolosan, France; INRAE, GeT-PlaGe, Genotoul, Castanet-Tolosan, France

## Abstract

The impact of host diversity on the genotypic and phenotypic evolution of broad-spectrum pathogens is a remaining issue. Here, we used populations of the plant pathogen *Ralstonia pseudosolanacearum* that were experimentally evolved on five types of host plants, either belonging to different botanical families or differing in their susceptibility or resistance to the pathogen. We investigated whether changes in transcriptomic profiles dissociated from genetic changes could occur during the process of host adaptation, and whether transcriptomic reprogramming was dependent on host type. Genomic and transcriptomic variations were established for 31 evolved clones that showed a better fitness in their experimental host than the ancestral clone. Few genomic polymorphisms were detected in these clones, but significant transcriptomic variations were observed, with a high number of differentially expressed genes (DEGs). In a very clear way, a group of genes belonging to the network of regulation of the bacterial virulence such as *efpR, efpH* or *hrpB*, among others, were deregulated in several independent evolutionary lineages and appeared to play a key role in the transcriptomic rewiring observed in evolved clones. A double hierarchical clustering based on the 400 top DEGs for each clone revealed two major patterns of gene deregulation that depend on host genotype, but not on host susceptibility or resistance to the pathogen. This work therefore highlights the existence of two major evolutionary paths that result in a significant reorganization of gene expression during adaptive evolution and underscore clusters of co-regulated genes associated to bacterial adaptation on different host lines.

**Data summary:** The authors confirm all supporting data, code and protocols have been provided within the article or through supplementary data files.

## Introduction

Understanding how pathogens adapt to their host or to new environmental conditions is still a major concern. Several studies have demonstrated the undeniable role of adaptive mutation fixation across generations to explain adaptation to changing environments or, in the case of pathogens, to new hosts [1, 2]. Here, we hypothesized that several strategies occur that will ultimately lead to the regulation of the same virulence and metabolism pathways and allow better adaptation of the pathogen.

In a previous work, we conducted an experimental evolution of the plant pathogen *Ralstonia pseudosolanacearum* strain GMI1000 through serial passage experiment into the stem of various plants [3]. The vast majority of the clones we obtained had significantly higher fitness gains than the ancestral clone. We then performed transcriptomic analyses of 10 clones evolved in the resistant tomato Hawaii 7996, which revealed a convergence towards a global rewiring of the virulence regulatory network [4]. As the evolution experiment was performed on several plant species [3], we sought to determine to what extent the transcriptional profiles mobilized for adaptation to tomato Hawaii were conserved for adaptation to other plants. We thus investigated genomic and transcriptomic variations in 21 additional clones obtained by experimental evolution of strain GMI1000 in either susceptible (tomato var. Marmande, eggplant var. Zebrina) and tolerant (bean var. blanc précoce, cabbage var. Bartolo) hosts.

## Methods

### Bacterial strains and growth conditions

The GMI1000 strain and a list of 21 evolved clones, derived from GMI1000, were investigated in this study (Table 1). The evolved clones were generated in our previous work by serial passage experiments (SPE) of the GMI1000 strain in four different plant species during 300 bacterial generations [3]. Five biological SPE replicates (named “A”, “B”, “C”, “D” and “E”) were conducted in parallel for each plant species thus generating five lineages of evolved clones per plant species (Table 1). For each evolved clone, the adaptive advantage was compared to the ancestral clone through the measure of their Competitive Index (CI) during *in planta* competition assays [3, 4]. The bacterial strains were revived from -80°C glycerol collections on agar plates containing BG medium supplemented with D-Glucose (5 g/l) and triphenyltetrazolium chloride (0.05 g/l) at 28°C [5]. For DNA and RNA extractions, bacterial strains were grown in MP synthetic liquid medium supplemented with L-Glutamine (10 mM) and oligoelements (1000 mg/l) at 28°C under agitation at 180 rpm [5]. The pH of the MP medium was adjusted to 6.5 with KOH. The composition of the oligoelements solution is given in supplemental file 1.

**Table 1.**
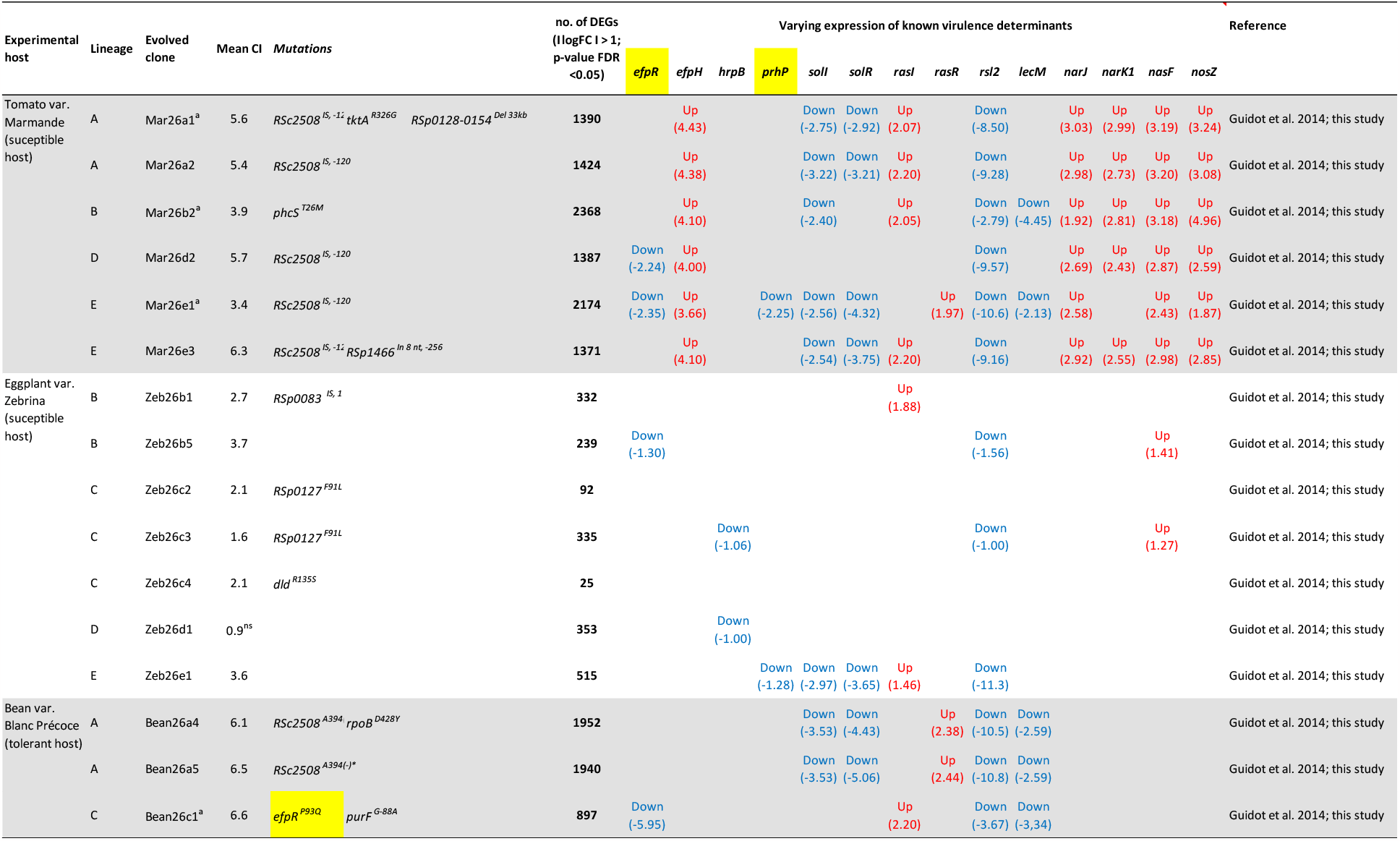
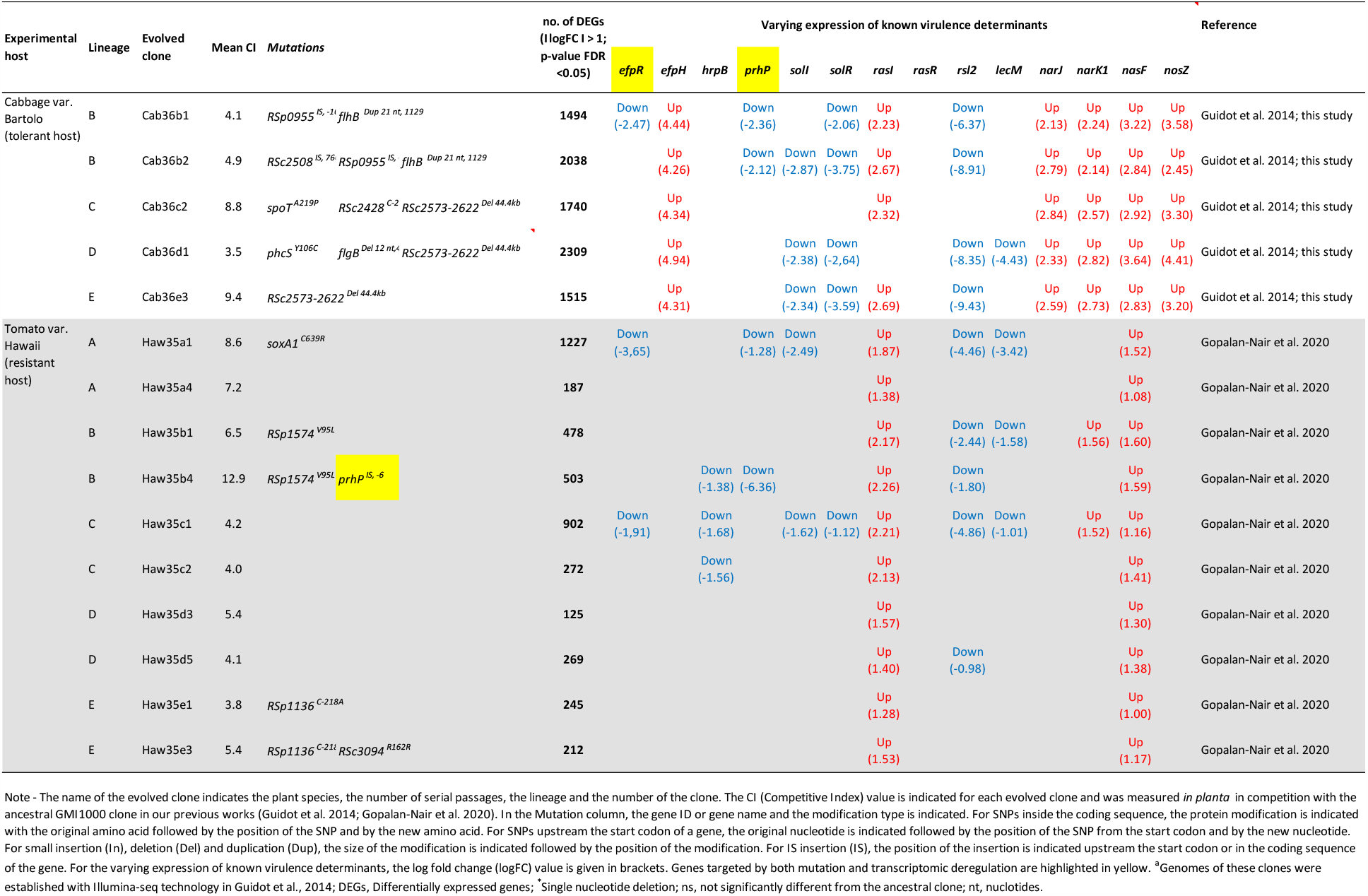
List of evolved clones with their genomic polymorphisms and transcriptomic variations of known virulence determinants.

### Genome sequencing and detection of genomic modifications

DNA were extracted from bacterial cultures in MP medium supplemented with 10 mM glutamine and collected at the beginning of stationary phase, in similar conditions as for Hawaii clones [4]. DNA sequencing was performed at the GeT-PlaGe core facility, INRAE Toulouse, France, and at Gentyane core facility, INRAE Clermont-Ferrand, France, using both the Illumina and PacBio sequencing technologies as for Hawaii clones [4].

Detection of genomic modifications was performed as described in our previous work [4]. All detected genomic modifications were checked by PCR amplification and sequencing with the Sanger technology using the primers reported in supplementary table S1.

### RNA extraction and RNA sequencing

RNA were extracted from bacterial cultures in MP medium supplemented with 10 mM glutamine and collected at the beginning of stationary phase, in similar conditions as for Hawaii clones [4]. Three biological replicates were conducted for each of the 21 clones and the GMI1000 ancestor strain. Oriented paired-end RNA sequencing (RNAseq) was performed on two lanes of an Illumina HiSeq3000 using a paired-end read length of 2×150 bp with the Illumina HiSeq3000 sequencing kits (Illumina) at the GeT-PlaGe core facility, INRAE Toulouse, France.

### Mapping and analysis of RNAseq data

Mapping and analysis of RNAseq data was performed as described in our previous work [4]. Differentially expressed genes were detected with EdgeR Bioconductor package version 3.30.3 [6]. Normalization was performed using TMM (trimmed mean of M-values) method [7].

Differentially expressed genes (DEGs) were called using the generalized linear model (GLM) likelihood ratio test using a False Discovery Rate (FDR) adjusted p-value < 0.05 [8]. Double hierarchical clustering analysis of the 400 top DEGs in the 31 evolved clones of *R. pseudosolanacearum* (21 clones from the present work and 10 clones from our previous work [4]) was performed using the hclust function from the stats R package (R Development Core Team, 2012) which computed Euclidean distances between clone profiles. Clustering was computed with Ward’s D2 method and a heat map was generated with the gplots R package.

## Results and discussion

We first investigated the genomic variations in the 21 clones compared to their ancestor by whole genome sequencing (Table 1). Both Illumina and Pacbio sequencing technologies were used for the detection of SNPs (Single Nucleotide Polymorphisms) or small InDels (Insertion-Deletion) and for the detection of large genomic rearrangements, respectively. These analyses revealed between zero and three genomic polymorphisms per clone compared to the ancestor. This low number of mutations is in the range of what we found in our previous sequenced evolved clones [3, 4] and included either SNPs, Indels, duplication, IS (Insertion sequence) insertion or large deletion (Table 1). All these genomic polymorphisms were confirmed by PCR amplification followed by Sanger sequencing or gel electrophoresis.

We then investigated the transcriptomic variations in the 21 evolved clones compared to their ancestor by RNA-sequencing approach. Analysis of RNA-seq data revealed that all samples rendered between 0.7 and 1.3 million of GMI1000-mapped reads. Differentially expressed genes (DEGs) between the evolved clones and the ancestral clone were considered as those presenting an FDR-adjusted *p*-value (padj,FDR) < 0.05 (Table 1 and Supplementary Table S2). In the 21 new investigated clones, when considering a log-fold change of expression I logFC I > 1, the number of DEGs varied between 25 and 2368 and was not correlated to the number of mutations (Table 1). A high number of DEGs was detected in the three clones evolved in eggplant Zebrina despite no detected genetic alteration, confirming a previous observation that some transcriptomic variations may be dependent on epigenetic modifications [4].

By looking more precisely at the DEGs, it appeared that there were groups of genes that were deregulated frequently, even recurrently, in several independent evolutionary lineages and regardless of the host considered (Table 1). In agreement with what was observed for the Hawaii clones, we detected a significant down-regulation for the virulence gene regulators *efpR, hrpB* and *prhP* [4] in 5, 2 and 4 of the newly 21 investigated clones, respectively. In addition, we observed a significant up-regulation for the *efpH* gene, a virulence regulatory gene homologous to *efpR* [9], but exclusively in clones evolved in tomato Marmande and in cabbage. We also observed significant variations in the expression pattern of genes involved in quorum-sensing-dependent virulence signaling pathways such as *solI/solR* [10] in 14 clones; *rasI*/*rasR* [11] in 24 clones; as well as the lectin encoding genes *lecM* and *rsl2* [12] in 21 clones. Genes involved in the denitrification pathway such as *narJ/narK1* and *nosZ* [13] were exclusively up-regulated in the tomato Marmande and cabbage clones and *nasF*, a nitrate transporter protein, was found up-regulated in 23 clones.

To go further in the analysis and increase its significance, we focused on the 400 top DEGs for each clone (200 most up-regulated and 200 most down-regulated). The comparison of these DEGs using a double hierarchical clustering is presented in Figure 1. This clustering showed that the evolved clones can be separated into two groups according to the host on which they have evolved, with two main branches clearly distinguishing the clones evolved on tomato Marmande and cabbage on the one hand and the clones evolved on tomato Hawaii, eggplant and bean on the other. This observation suggested that there are at least two major patterns of gene deregulation associated with strain GMI1000 adaptation to its hosts, and that these patterns are associated to the host genotype but not to the host susceptibility to bacterial wilt disease. In terms of discriminating genes, it appears that the Tomato Marmande / cabbage branch is associated with an upregulated expression of cluster (1) (Figure 1 and Supplementary Table S2), which includes 180 genes, among which *efpH, narJ* and *nosZ*. The analysis also confirmed down-regulation of part of the *hrpB* regulon (cluster 4) in several clones evolved on Tomato Hawaii, eggplant and Bean, showing significant differences in expression of several genes encoding type 3 effectors in Hawaii evolved clones [4].

**Figure 1.**
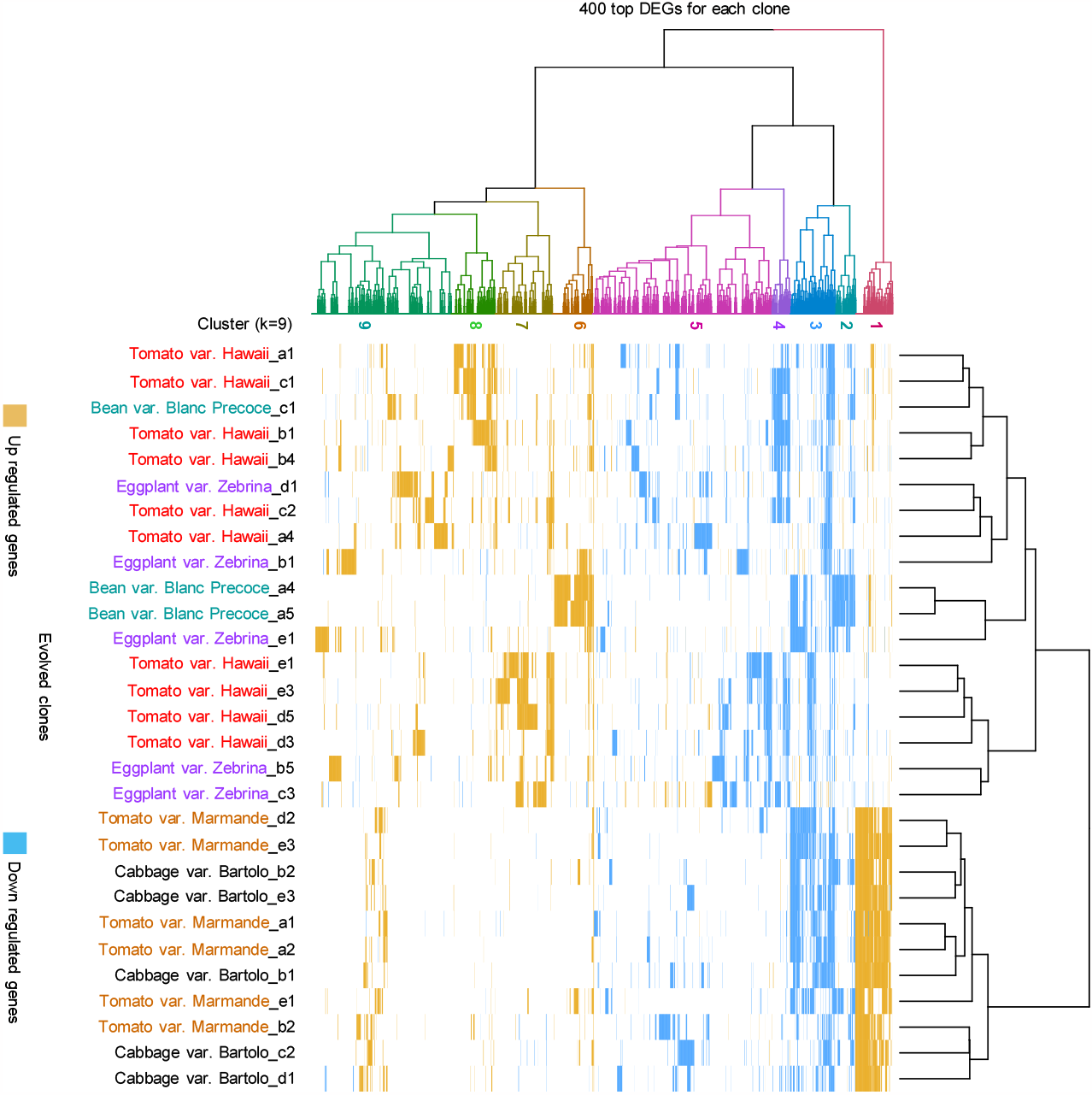
Double hierarchical clustering analysis of the 400 top Differentially Expressed Genes (DEGs) in 31 evolved clones of *R. pseudosolanacearum*. The 400 top DEGs (FDR<0.05) (200 most up-regulated and 200 most down-regulated) were selected for each clone. Down- and up-regulated genes were binary coded, respectively -1 (blue) and 1(orange). The hclust function from the stats R package (R Development Core Team, 2012) computed Euclidean distances between clone profiles. Clustering was computed with Ward’s D2 method and a heat map was generated with the gplots R package

Altogether, these data highlight a significant reorganization of gene expression associated to adaptation of *R. pseudosolanacearum* to multihost species, which converged toward two major patterns of gene deregulation according to the host genotype. In all cases, a relatively small number of genes (including several transcriptional regulators such as *efpR, efpH, hrpB, rasI*/*R, solI*/*R, rsl2*) seem to play a key role in these transcriptomic rewiring. The distribution of these regulatory genes in different clusters of co-regulated genes opens avenues for further characterization of these regulons and possible cross-regulations associated to adaptive process to host plants.

## Supplementary material

**Supplementary table S1** List of primers used in this study

**Supplementary Table S2** RNAseq raw data and analysis for all wild-type (wt) GMI1000 strain and all 31 experimentally evolved clones transcripts detected in minimal medium supplemented with 10mM glutamine. The 400 top DEGs identified for each clone and used for the double hierarchical clustering analysis are indicated by a cross in the ‘Top DEGs’ column. For each clone, the top DEGs are encoded -1.00 for the down-regulated genes, 1.00 for the up-regulated genes and 0.00 for the genes not differentially regulated compared to the wt strain. The gene cluster to which the 400 top DEGs in each 31 evolved clone belong is given in the ‘cluster (k=9)’ column. RNAseq data analysis for the Zeb26c2 and Zeb26c4 clones are given in grey at the end of the table. The number of DEGs for these two clones were not sufficient to be included in the double hierarchical clustering analysis.

**Supplemental file 1** Composition of the oligoelements solution 1000X (250 ml) and protocol

