## Supplemental File 1 for "Transcriptomic profiling reveals host-specific evolutionary pathways promoting enhanced fitness in the plant pathogen *Ralstonia pseudosolanacearum*"

**Gopalan-Nair et al.**

**Supplemental file 1**

### Composition of the oligoelements solution 1000X (250 ml) and protocol

This oligoelement base composition is derived from the Hutner's trace element. The protocol should provide a stable solution, i.e. no salt precipitation, over 4°C storage.

*1. Iron solution.*

For 250 ml, dissolve the listed salts in 100 ml of distilled water in the order indicated:

| | |
|---|---|
| $FeSO_4$, 7 $H_2O$ | 1.25 g |
| $Na_2EDTA$, 2 $H_2O$ | 12.50 g |

Adjust the pH with KOH 10N until complete dissolution of EDTA and having a golden yellow solution that should be around pH 8.

*2. Trace solution.*

Dissolve the listed salts in 100 ml of distilled water in the order indicated:

| | |
|---|---|
| $ZnSO_4$, 7 $H_2O$ | 5.50 g |
| $H_3BO_3$ | 2.85 g |
| $MnCl_2$, 4 $H_2O$ | 1.26 g |

| | |
|---|---|
| $CoCl_2$, 6 $H_2O$ | 0.40 g |
| $CuSO_4$, 5 $H_2O$ | 0.39 g |
| $(NH_4)_6Mo_7O_{24}$, 4 $H_2O$ | 0.28 g |

*3. Combining 1 + 2 solutions.*

Combine the solution 1 and the solution 2, and readjust the pH to 6.5 using 10N KOH.

Bring the final volume to 250 ml with distilled water.

Sterilize the solution by filtering using 0.22 μm filter.

Store at 4°C.

The Oligo solution is initially bright green, turning purple upon storage. Precipitates should never be formed.
